# The Emerging Role of 3-Hydroxyanthranilic Acid on *C. elegans* Aging Immune Function

**DOI:** 10.1101/2024.01.07.574394

**Authors:** Luis S. Espejo, Destiny DeNicola, Leah M. Chang, Vanessa Hofschneider, Anne E. Haskins, Jonah Balsa, Samuel S. Freitas, Angelo Antenor, Sage Hamming, Bradford Hull, Raul Castro-Portuguez, Hope Dang, George L. Sutphin

**Affiliations:** Molecular & Cellular Biology, University of Arizona, Tucson, AZ, USA

## Abstract

3-hydroxyanthranilic acid (3HAA) is considered to be a fleeting metabolic intermediate along tryptophan catabolism through the kynurenine pathway. 3HAA and the rest of the kynurenine pathway have been linked to immune response in mammals yet whether it is detrimental or advantageous is a point of contention. Recently we have shown that accumulation of this metabolite, either through supplementation or prevention of its degradation, extends healthy lifespan in *C. elegans* and mice, while the mechanism remained unknown. Utilizing *C. elegans* as a model we investigate how 3HAA and *haao-1* inhibition impact the host and the potential pathogens. What we find is that 3HAA improves host immune function with aging and serves as an antimicrobial against gram-negative bacteria. Regulation of 3HAA’s antimicrobial activity is accomplished via tissue separation. 3HAA is synthesized in the *C. elegans* hypodermal tissue, localized to the site of pathogen interaction within the gut granules, and degraded in the neuronal cells. This tissue separation creates a new possible function for 3HAA that may give insight to a larger evolutionarily conserved function within the immune response.

**Graphical Abstract:** 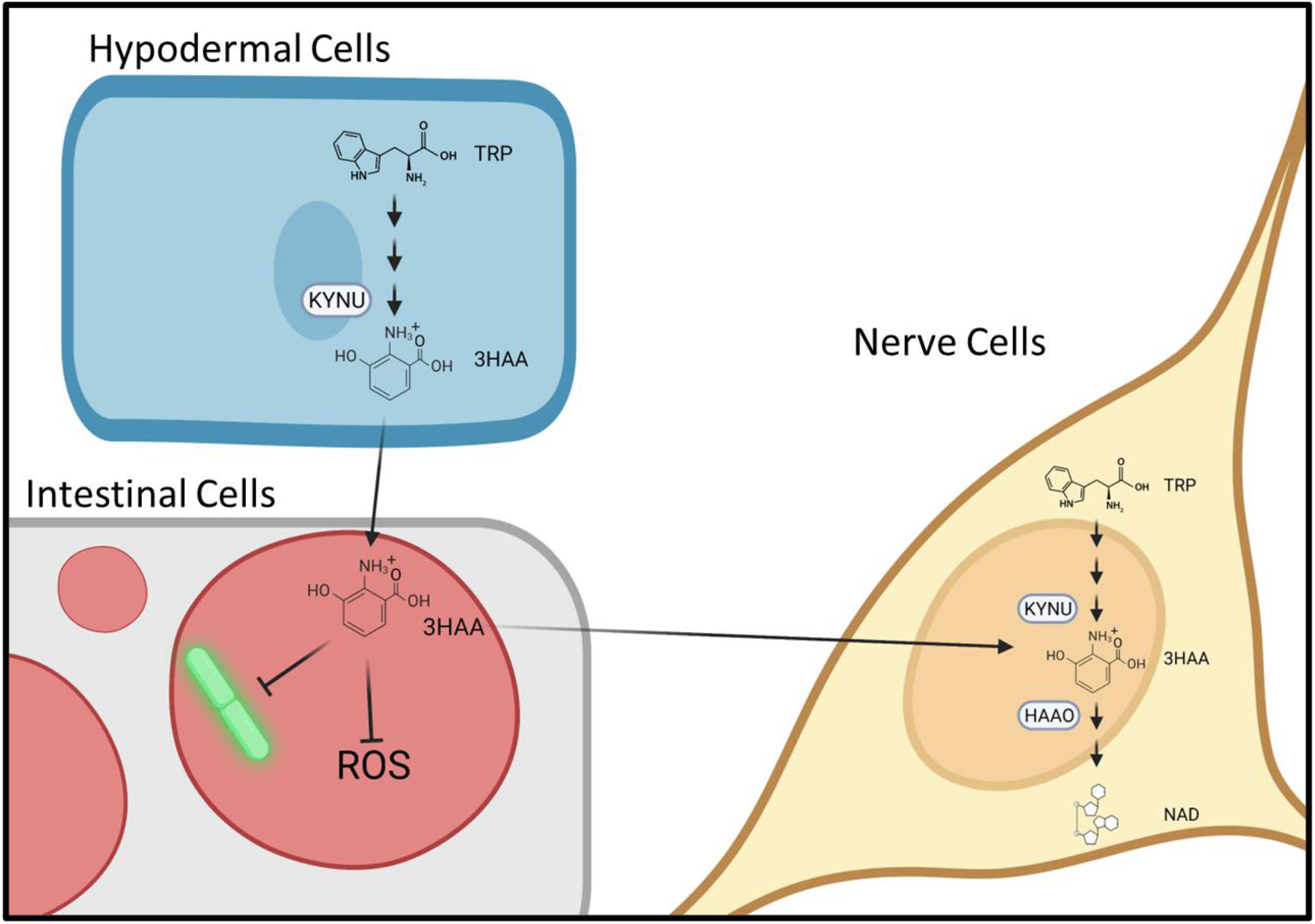

## Introduction

*Caenorhabditis elegans* has been a central model in diverse areas of age-related research since the 1980s^1^, and has served as a model for innate immunity and host-pathogen interaction since the early 1990s.^2–4^ *C. elegans* have a set number of somatic cells in adulthood but recapitulate aspects of molecular immune signaling which are orthologous to humans.^5–8^ We previously found that accumulation of the tryptophan metabolite 3-hydroxyanthranilic acid (3HAA) via supplementation or inhibition of its metabolizing enzyme 3-hydroxyanthranilate 3,4-dioxygenase (HAAO) increases lifespan and extends many aspects of healthy aging in *C. elegans* and murine models.^9^ 3HAA and other kynurenine pathway metabolites have been characterized as both pro-inflammatory and pro-immunogenic as well as anti-inflammatory and immunosuppressive.^10–21^ A similar conflicting dichotomy is seen in the kynurenine pathway metabolites in regards to being pro-oxidant and antioxidant.^22–27^ To evaluate therapeutic potential of 3HAA and kynurenine pathway manipulations, it is important to understand precisely what impact they have on aging immune health.

The dual natures of 3HAA argues for a context specific application of the metabolite, making the synthesis, localization, and degradation important factors in understanding the impact of the metabolite on immune function. For example the ability of 3HAA to be pro or anti-oxidant appears to be intricately linked to environmental factors like metal ions.^25^ 3HAA has also been shown to reduce essential metal ions such as iron.^28^ Our initial findings demonstrate that 3HAA increases markers associated with anti-inflammatory responses as well as effectively neutralizes H2O2 in vitro.^9^ These findings, among others, suggest an inhibitory role of 3HAA on inflammation and immune response but the overall impact on the immune function and the ability to combat pathogens during the aging process in undefined.^12^

Here we utilized *C. elegans* as a model to investigate the impact that elevated 3HAA has on aging immune function. Animals lacking *haao-1*, the gene encoding HAAO in *C. elegans*, are resistant to age-associated immune decline and better handle pathogenic challenge. *haao-1* mutants experienced lower rates of *Escherichia coli* infection and live longer once infected than their wild counterparts. Compared to the weaker pathogen *E. coli*, the kynurenine pathway is upregulated to a greater extent in response to the more pathogenic gram-negative *Pseudomonas aeruginosa*. In the absence of *haao-1* animals accumulate 3HAA with age, which may in turn be consumed in the defense against pathogens. The immune benefits of *haao-1* inhibition do not appear to operate through canonical *C. elegans* immune signaling pathway, but rather through direct antimicrobial activity of 3HAA itself, which is selectively localized to sites of pathogen interface within the intestinal enterocytes. This research provides a novel mechanistic role for 3HAA in *C. elegans* innate immune function and pathogen resistance.

## Results

The kynurenine pathway catabolizes tryptophan(**Fig. 1A**) and is responsible for the majority of tryptophan metabolism in mammals.^29^ It also is responsible for the biosynthesis of NAD+ contributing to the NAD+ pool along with dietary intake. We previously found that inhibition of *haao-1* extends lifespan in *C. elegans* mediated by physiological accumulation of the HAAO substrate 3HAA. The red 3HAA becomes visible under a dissection microscope by 11 days of age, or day 8 of adulthood (**Fig. 1B**)^9^. *C. elegans*, like mammals, experience a decline in immune function that results in increased susceptibility to pathogenic challenge with age.^30^ Given the established role of kynurenine metabolism in immune function in mammals,^31,32^ we asked whether kynurenine pathway interventions that alter lifespan would similarly impact immune function in *C. elegans* during aging. *C. elegans* with deletion mutations in four genes—*kmo-1, kynu-1, haao-1*, and *umps-1*—were raised on a live gram-negative *E. coli* food source (strain HT115, which are mildly pathogenic to *C. elegans*)^33^ and challenged with the highly pathogenic gram-negative PA14 strain *P. aeruginosa* at the 4^th^ larval stage (L4). Deletion of *kynu-1* and *haao-1*, which encode kynureninase (KYNU) and HAAO, the enzymes are responsible for the synthesis and degradation of 3HAA, respectively and are both long lived on *E. coli*^9,34^ and modestly resistant to PA14 challenge at L4 (**Fig. 1C**).

**Figure 1:**
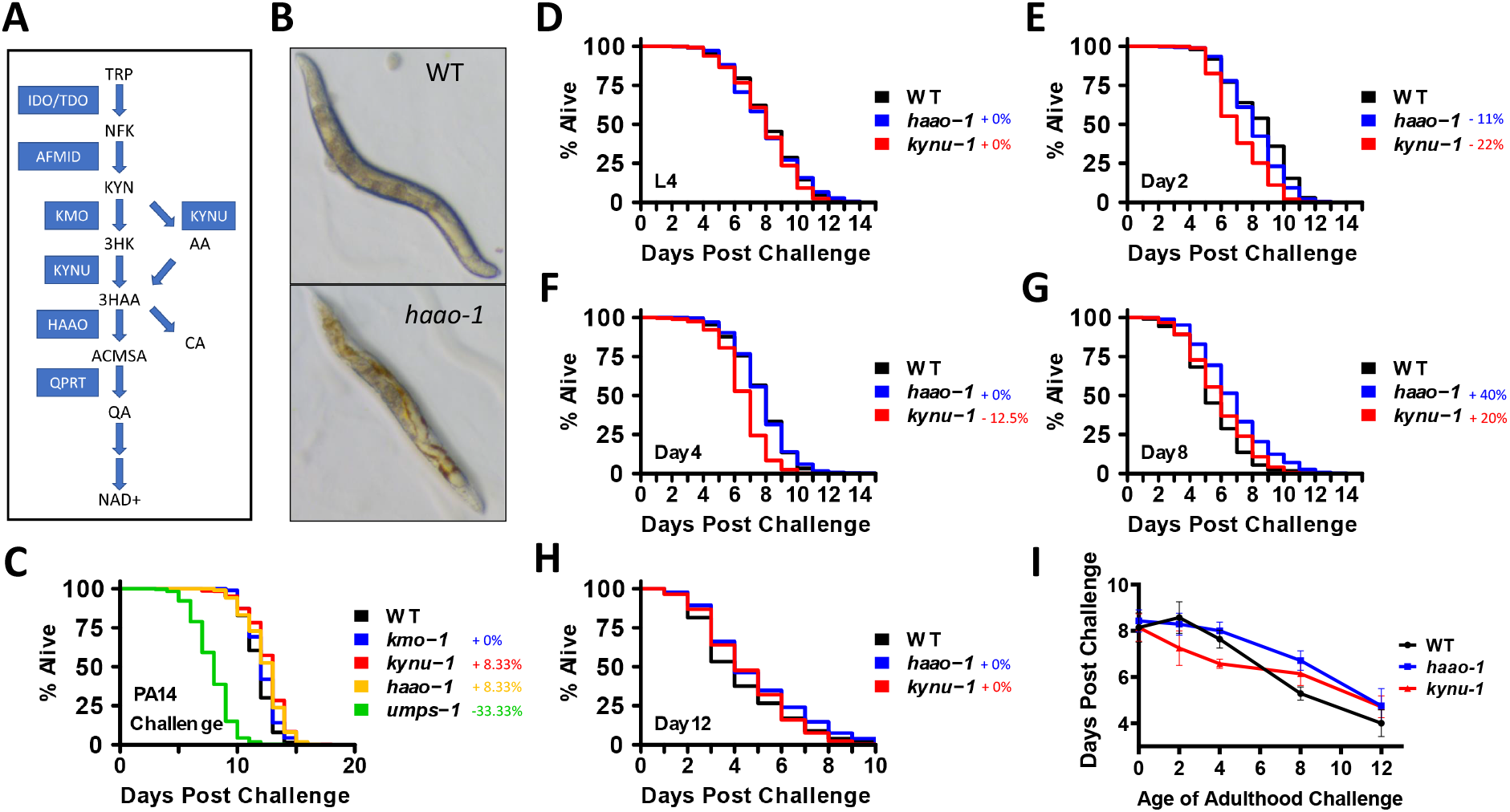
*haao-1* Mutants Resistant to Age-Associated Immune Decline. **(A)** The kynurenine pathway is responsible for a majority of the metabolism of tryptophan and the endogenous production of NAD+. **(B)** *haao-1* mutants grow visibly red with age due to the accumulation of the 3HAA metabolite. **(C)** Kynurenine pathway mutants synchronized on HT115 EV RNAi differentially impact host survival upon *P. aeruginosa* challenge at L4 stage. **(D)** Survival of haao-1 and kynu-1 mutants synchronized on OP50 challenge with *P. aeruginosa* at L4 stage. **(E-H)** C. elegans survival reared on OP50 bacteria then challenged with *P. aeruginosa* at days: 2, 4, 8, and 12 of adulthood. **(I)** Characterization of age-associated immune decline measuring median survival post *P. aeruginosa* challenge.

To investigate how *kynu-1* and *haao-1* impact *C. elegans* pathogen resistance during aging, *kynu-1* and *haao-1* animals were raised on *E. coli* (strain OP50, which is slightly more pathogenic that HT115) and challenged with PA14 at different ages (**Fig. 1D-H**). *kynu-1* knockout animals displayed a decreased pathogen resistance during early adulthood (**Fig. 1G**). In contrast, *haao-1* knockout animals had similar pathogen resistance to wild type during early adulthood, but were more resistant than wild type when challenged at later ages (**Fig. 1I**). These results indicate that *haao-1* deficient animals do not experience the same level age-associated immune decline or *kynu-1* mutants.

*C. elegans* have several well-established immune signaling pathways. To test whether *haao-1* inhibition improves immune response via these established pathways, epistasis survival experiments were conducted (**Fig. 2A-C**). Lifespan extension via *haao-1* inhibition was not rescued by *daf-16* (**Fig 2A**), *pmk-1* (**Fig 2B**), or *dbl-1* (**Fig. 2C**) RNAi, indicating that the age-associated benefits form *haao-1* inhibition is likely not acting through classical *C. elegans* immune signaling pathways. In *haao-1* deficient animals, 3HAA is consistently elevated relative the wild type during aging. This elevation appears to modestly increase over time through day 4 of adulthood, and then increase dramatically between day 4 and 8 of adulthood (**Fig. 2D**). During the aging process, the ratio of kynurenine (KYN) to tryptophan (TRP), a common measure of kynurenine pathway activation that is used as a measure of IDO/TDO activity, decreases in wild type, *kynu-1* and *haao-1* mutants (**Fig. 2E**). *kynu-1* mutants, and to a lesser extend *haao-1* mutants, have a reduced KYN:TRP ratio throughout lifespan, which is likely due to elevated KYN levels (**Fig. S3**) resulting from impaired KYN degradation, rather than higher activity of TDO, as tryptophan is not reduced in these backgrounds (**Fig. S3**).

**Figure 2.**
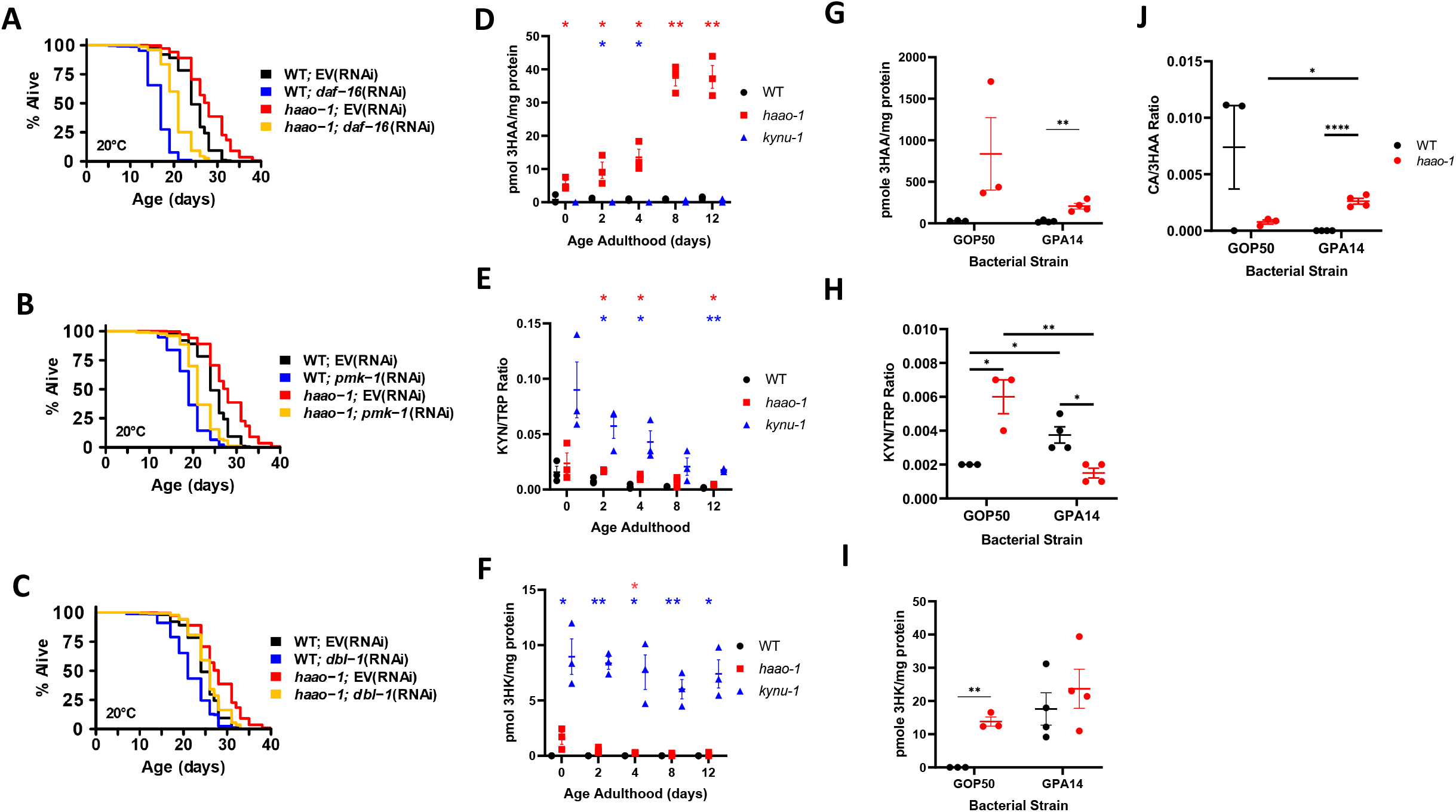
Immune genes and Metabolic Activity. **haao-1 Mutant Improved Immune Function is Independent of Classical C. elegans Immune Signaling Funneling Pathway Flux Towards 3HAA. (A)** Epistatic survival of *haao-1* mutants using *daf-16* HT115 knock down. **(B)** Epistatic survival of *haao-1* mutants using *pmk-1* HT115 knock down. **(C)** Epistatic survival of haao-1 mutants using *dbl-1* HTT115 knock down. **(D)** Mass spectrometry of 3HAA metabolite over course of aging normalized to total population protein. **(E)** KYN/TRP ratio throughout aging up to day 12 of adulthood, classically used to represent flux through the kynurenine pathway. **(F)** Mass spectrometry of 3HK metabolite throughout aging normalized to total population protein. **(G)** Mass spectrometry of 3HAA metabolite of *haao-1* mutants and WT animals under low and high pathogenic conditions of GOP50 and GPA14. **(H)** KYN/TRP ratio of *haao-1* and WT under different pathogenic challenges obtained via mass spectrometry. **(I)** Mass spectrometry of 3HK metabolite in *haao-1* and WT under different pathogenic challenges. **(J)** CA/3HAA metabolite ratio determined via mass spec representing oxidation of 3HAA to CA.

We next examined how kynurenine pathway metabolites change when wild type or *haao-1* knockout worms are exposed to mildly pathogenic (*E. coli* strain OP50) vs. highly pathogenic (*P. aeruginosa* strain PA14) bacteria. In wild type animals, the KYN:TRP ratio increases following exposure to PA14, driven by a decrease in TRP and an increase in KYN (Fig. S2), suggesting activation of TDO-2 to increase flux through the kynurenine pathway (**Fig. 2H**). This indicates that *C. elegans* promote kynurenine pathway flux via activation of TDO-2 in the presence of strong pathogens, paralleling IDO activation in mammals.^35^ The opposite relationship is seen in the *haao-1* mutants, with the KYN:TRP ratio elevated relative to wild type animals on *E. coli*, but strongly repressed following PA14 challenge (**Fig. 2H**). This flip in KYN/TRP in *haao-1* mutants is likely due to *haao-1* mutants accumulating KYN in low pathogen environments, potentially due to feedback inhibition of KYNU, but tryptophan levels follow the same trend as wild type animals (**Supplemental Fig. 2**). Additionally, there is a trend toward increased in 3-hydroxykynurenine (3HK), the metabolite that is converted to 3HAA by KYNU (**Fig. 1A**), in wild type animals under strong pathogenic challenge (**Fig. 2I**) suggesting an increased flow towards 3HAA during challenge. When animals age on the standard OP50 diet 3HK levels decrease over time (**Fig. 2F**). While 3HAA itself decreases in response to PA14 challenge (**Fig. 2G**), the ratio of the oxidized product of 3HAA, cinnabarinic acid (CA), to 3HAA increases **(Fig. 2J**) indicating potential consumption of 3HAA in defense against strong pathogens. Taken together, this evidence suggests the kynurenine pathway is upregulated under strong pathogenic challenge and that 3HAA is consumed in the process of host defense.

To understand how *haao-1* inhibition and 3HAA impact the host-pathogen relationship, we examined the impact *haao-1* deletion on the dynamics of bacterial infection using a new system that we recently developed called Systematic Imaging of *Caenorhabditis* Killing Organisms (SICKO)^36^. SICKO allows several metrics of infection with fluorescently-labeled bacteria to be monitored over time in individual free-crawling *C. elegans*^36^, including infection initiation, progression, and subsequent survival. We exposed wild type and *haao-1* knockout worms to *E. coli* OP50 constitutive expressing GFP for 7 days, transferred randomly selected animals to single worm culture environments^37^, and monitored subsequent infection dynamics and survival over time. Both *haao-1* mutants and wild type animals experienced death associated with infection, but *haao-1* mutants experience a decreased rate of infection and infection associated death (**Fig. 3A**). Additionally, wild animals experienced an increase onset of infection post challenge (**Fig 3B**). Infected *haao-1* knockout animals performed significantly worse than their non-infected counterparts indicating toxicity of GOP50 is less impactful than infection in relation to death. Once infected, *haao-1* animals have improved survival relative to wild type (**Fig. 3C**). SICKO calculates a infection index (aka the SICKO Score) that incorporates information about current level of infection in live animals and the number of past deaths in infected animals, providing a composite measure of infection severity that can be used to compare populations with distinct survival and infection dynamics over time^36^. *haao-1* mutants experienced lower infection index relative to wild type animals throughout life (**Fig. 3D**).

**Figure 3.**
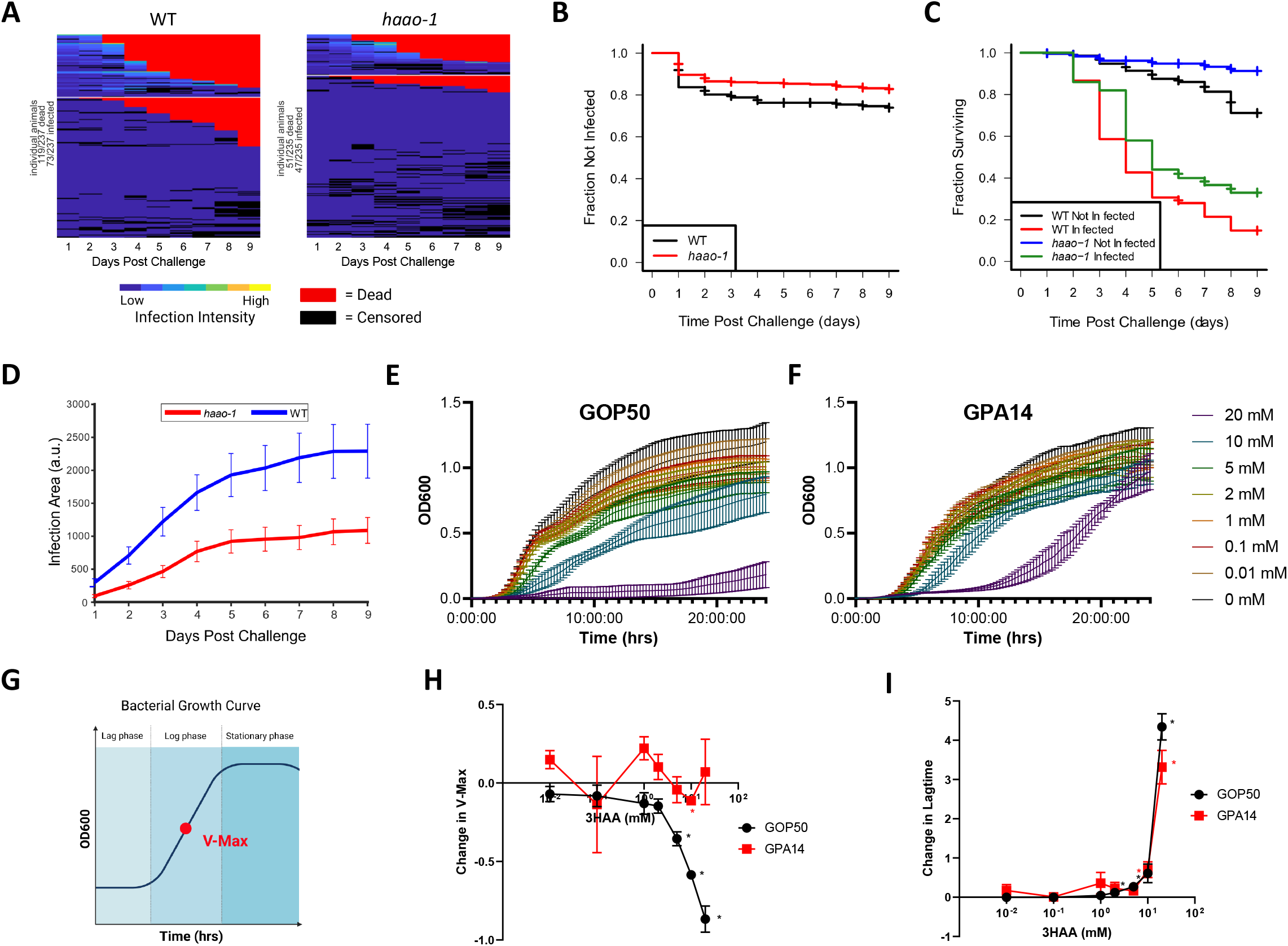
SICKO and Bacteria Response to 3HAA. ***haao-1* Mutants have Improved Response to Pathogens while the Accumulating 3HAA has Strong Antimicrobial Activity. (A)** SICKO (Systematic Imaging of *Caenorhabditis* Killing Organisms) infection tracking of GOP50 infection area in WT and *haao-1* animals after 7-day pathogenic challenge. All animals above the white line are infected in each condition. WT animals were significantly more associated with infection (p < 0.005; chi-squre). In both genetic backgrounds death was associated with infection (p < 0.05; chi-square). **(B)** Onset of detectable infections in WT and *haao-1* animals using SICKO system following pathogenic challenge. *haao-1* animals had significantly different onset of infection (p < 0.05, log-rank test). (C) Survival

These findings indicate that HAAO inhibition is sufficient to improve host response to pathogens in terms of resistance to both infection initiation and progression. Because 3HAA is elevated in *haao-1* knockout animals, is consumed during PA14 infection, and mimics lifespan extension from *haao-1* knockdown, and because lifespan extension from *haao-1* knockout is not dependent on canonical immune pathways, we next asked whether 3HAA might media the improved pathogen resistance of *haao-1* deficient animals by directly impacting bacteria survival and growth. In both GOP50 and GPA14 cultured in liquid media, 3HAA inhibited bacterial growth in a dose-dependent manner (**Fig. 3E-F**). Different aspects of bacterial growth dynamics can be quantified from the bacterial growth curves; V-max provides insight to maximum growth rate of the bacteria within the medium, while lag time is a reflection of the size of the initial population and is thus an indication of initial impact of an intervention on bacterial death. 3HAA inhibits the GOP50 V-max in a dose-dependent manner (**Fig. 3G**). Here we found that both pathogen lagtimes directly correlate to bacteria concentration (Supplemental Fig. 4). The lagtime for both the weak and strong gram-negative pathogens is significantly impacted by 3HAA in a dose dependent manner (**Fig. 3I**) indicating 3HAA prevents initiation of bacterial growth in early stages or kills a subset of the bacteria on first exposure.

**Figure 4:**
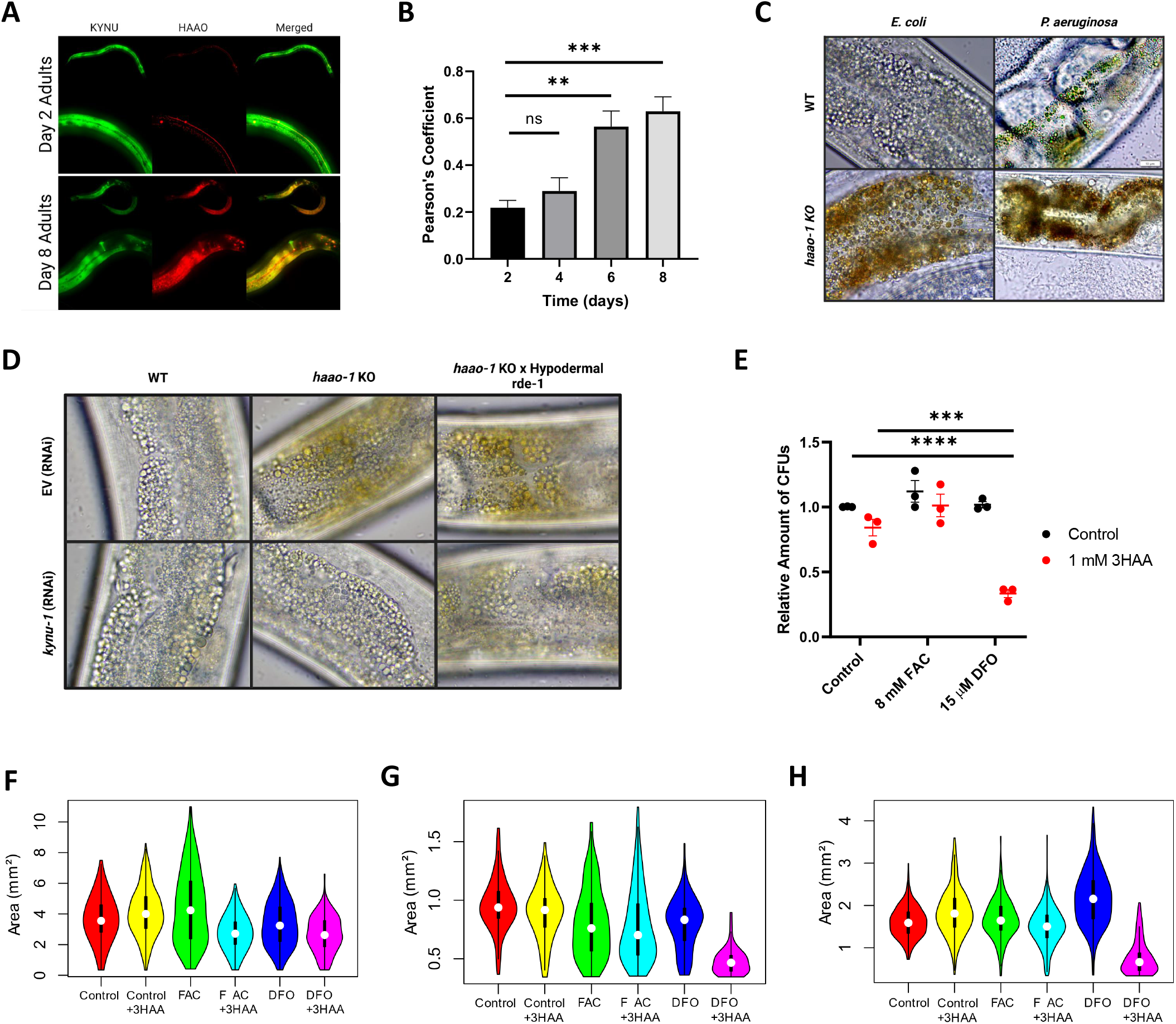
3HAA Displays Special Temporal Separation and Antimicrobial Activity. **(A)** 3HAA synthesizing enzyme KYNU-1::GFP and 3HAA degrading enzyme HAAO-1::mScarlet at day 2 and day 8 of adulthood. **(B)** Pearson coefficient of KYNU-1::GFP and HAAO-1:mScarlet in 3-dimensions of entire animal at days 2, 4, 6, and 8 of adulthood. **(C)** Color 40X microscopy of C. elegans gut epithelium directly below the pharyngeal pump when challenged with OP50 and PA14. **(D)** WT, *haao-1* mutants, and *haao-1* mutants crossed with selectively active *rde-1* (enabling RNAi knock-down only in the hypodermal tissue), exposed to EV and *kynu-1* RNAi until day 6 of adulthood. **(E)** Count of GOP50 CFUs relative to control LB agar plates with ammonium ferric citrate (FAC) for iron supplementation and deferoxamine (DFO) for iron Chelation. **(F-H)** Area of GOP50 **(F)**, GPA14 **(G)**, and GHT115 **(H)** CFUs on LB agar plates when supplemented with 1 mM 3HAA, 8 mM FAC, and 15 μM DFO. White dots represent mean while black bars indicate SD. GOP50 on DFO + 3HAA were 27% (p < 0.0005; t-test) smaller while GPA and GHT115 on DFO + 3HAA were roughly 50% smaller (p < 0.0005; t-test)

To understand what context 3HAA may potentially contact pathogens it is critical to pinpoint its location, site of synthesis, and site of degradation. Early in adulthood animals display a distinct separation of 3HAA synthesis and degradation (**Fig. 4A-B**). Synthesis of 3HAA via KYNU-1 appears to happen across many tissues of the animal and heavily in the hypodermis. While degradation via HAAO-1 only occurs in a small subsection of tissue in which synthesis occurs. HAAO-1 during early adulthood primarily is located within the neurons (Supplemental Fig. 5). Surprisingly, this special temporal separation of synthesis and degradation becomes lost as the animal passes 6 days of adulthood (**Fig. 4B**). This special temporal gap creates a expands the potential localization of 3HAA between the process of metabolism through the kynurenine pathway.

Upon further inspection of 3HAA accumulation in haao-1 mutants, 3HAA appears not to be at the site of synthesis or degradation but rather located in gut granule structures within the *C. elegans* intestine (**Fig. 4C**). Due to the green pigmentation of PA14 toxins (Supplemental Fig. 5) animals challenged with the pathogen can also be observed with remnants of the pathogen in similar gut granular structures. Additionally, *haao-1* mutants challenged with PA14 have no distinctly separate green or red granular structures, indicating that 3HAA may contact pathogens within the subcellular structure. This finding potentially uncovers a previously unknown immune role and regulation of 3HAA in *C. elegans* pathogen combat. To distinguish if the 3HAA in the gut granules is from a distal tissue a *rde-1*, required for RNAi impact, mutant selectively functional in the hypodermis was utilized. Indeed, when kynu-1 is inhibited in the hypodermis, significant amounts of 3HAA fail to accumulate in the gut granules (**Fig. 4D**). Taken together this demonstrates that 3HAA is synthesized in a distal tissue and is transported to the pathogen interface.

The *C. elegans* gut granules have been proposed to be heavily regulated in regard to metal ion availability. 3HAA is also known to interact with iron leaving the potential of synergistic interactions in vivo. To test this possibility 3HAA was combined with iron perturbations to assess solid surface growth of pathogens. A strong synergistic impact is seen between 3HAA and the iron chelator deferoxamine (DFO) (**Fig. 4E**). DFO on it’s own does not have a strong impact on bacterial growth in solid or liquid culture (**Fig. S6**). Not only I does the combination of DFO and 3HAA inhibit bacterial ability to form colonies, the colonies capable of growing are significantly smaller than control colonies, or DFO and 3HAA alone (**Fig. 4F-G**). 1mM 3HAA by itself does not significantly impact bacterial growth but combined with likely metal perturbations in gut granules accounts for a potent antimicrobial activity of a conserved immune response in *C. elegans*.

## Discussion

The kynurenine is evolutionarily conserved and has been tied to immune function in mammals with contradictory indications to its role. We previously demonstrated that *haao-1* inhibition and 3HAA supplementation was capable of extending healthy lifespan in *C. elegans* and mice. Here we see that kynurenine pathway activity is linked to *C. elegans* immune function. In particular *haao-1* inhibition and 3HAA serve to decrease age-associated immune decline and help the host combat pathogen. The pathway is activated in response to strong pathogenic challenge and flux is funneled toward the production of 3HAA which is consumed in the process of combating pathogen. *haao-1* animals also performed better under pathogenic challenge. They survived stronger pathogenic challenges with age in addition to fending off weaker pathogenic challenges and slowing the progression of infection throughout the population.

Surprisingly, it does not appear that *haao-1* effects on *C. elegans* immune function occur through the canonical *C. elegans* immune signaling pathways. It appears that the major activity of 3HAA occurs on the pathogen itself. This activity is regulated by localization of 3HAA to the gut cell epithelium. It seems likely that the 3HAA resides in the gut granules because another kynurenine metabolite, anthranilic acid, is also present in that same sub cellular structure.^38,39^ In the gut granules, when 3HAA comes into contact with pathogens its antimicrobial activity would may serve as a host defense against invading pathogens. Regulation of antimicrobial activity is only possible via tissue separation in synthesis and degradation. By being synthesized in the hypodermis, localized in the gut granules, and degraded in the neurons, 3HAA can reach higher concentrations than if it was distributed throughout the body. This may explain why 3HAA has been viewed as a fleeting metabolic intermediate in the past; as if one only looks at sites where it is manufactured or degraded the 3HAA itself may not be in abundant levels. Additionally, antimicrobial affects of 3HAA may be intricately linked to metal ion availability. 3HAA has been noted to interact with ions in the past acting as a reducing agent. This would not be the first account of intermediate function as 3HAA has been proposed as a microbial attack mechanism in conjunction with iron sequestration in *Cryptococcus neoformans* and other organisms.^40,41^ Because elevated 3HAA and kynurenine pathway activity has been witnessed in patients undergoing immune challenge this research may provide insight into 3HAA’s evolutionary role in immune function. 3HAA’s elevation may due to immune response mounting against potential pathogens. This impact of 3HAA may also be context specific. In one tissue it may combat pathogens while in another may assist in oxidative stress. This creates multiple benefits that all may stem from the conserved metabolite.

## Acknowledgements

This work was made possible by the following funding: R35GM133588, T32GM008659, T32AG058503.

## References

1. Johnson, T. E. & Wood, W. B. Genetic analysis of life-span in Caenorhabditis elegans. Proc. Natl. Acad. Sci. U. S. A. 79, 6603–6607 (1982).

2. Pukkila-Worley, R. & Ausubel, F. M. Immune defense mechanisms in the Caenorhabditis elegans intestinal epithelium. Curr. Opin. Immunol. 24, 3–9 (2012).

3. Taffoni, C. & Pujol, N. Mechanisms of innate immunity in C. elegans epidermis. Tissue Barriers 3, e1078432 (2015).

4. Pujol, N. et al. Distinct innate immune responses to infection and wounding in the C. elegans epidermis. Curr. Biol. CB 18, 481–489 (2008).

5. Kurz, C. L. & Tan, M.-W. Regulation of aging and innate immunity in C. elegans. Aging Cell 3, 185–193 (2004).

6. Marsh, E. K. & May, R. C. Caenorhabditis elegans, a Model Organism for Investigating Immunity. Appl. Environ. Microbiol. 78, 2075–2081 (2012).

7. Arras, L. D. et al. An Evolutionarily Conserved Innate Immunity Protein Interaction Network. J. Biol. Chem. 288, 1967–1978 (2013).

8. Elkabti, A. B., Issi, L. & Rao, R. P. Caenorhabditis elegans as a Model Host to Monitor the Candida Infection Processes. J. Fungi 4, (2018).

9. Dang, H. et al. 3-hydroxyanthranilic acid – a new metabolite for healthy lifespan extension. 2021.06.01.446651 Preprint at 10.1101/2021.06.01.446651 (2021).

10. Lowe, M. M. et al. Identification of Cinnabarinic Acid as a Novel Endogenous Aryl Hydrocarbon Receptor Ligand That Drives IL-22 Production. PLoS ONE 9, e87877 (2014).

11. Hoshi, M. et al. 3-Hydroxykynurenine Regulates Lipopolysaccharide-Stimulated IL-6 Production and Protects against Endotoxic Shock in Mice. ImmunoHorizons 5, 523–534 (2021).

12. Christen, S., Thomas, S. R., Garner, B. & Stocker, R. Inhibition by interferon-gamma of human mononuclear cell-mediated low density lipoprotein oxidation. Participation of tryptophan metabolism along the kynurenine pathway. J. Clin. Invest. 93, 2149–2158 (1994).

13. Liu, X., Newton, R. C., Friedman, S. M. & Scherle, P. A. Indoleamine 2,3-dioxygenase, an emerging target for anti-cancer therapy. Curr. Cancer Drug Targets 9, 938–952 (2009).

14. Sekine, H. et al. Hypersensitivity of Aryl Hydrocarbon Receptor-Deficient Mice to Lipopolysaccharide-Induced Septic Shock. Mol. Cell. Biol. 29, 6391–6400 (2009).

15. Huttunen, R. et al. HIGH ACTIVITY OF INDOLEAMINE 2,3 DIOXYGENASE ENZYME PREDICTS DISEASE SEVERITY AND CASE FATALITY IN BACTEREMIC PATIENTS. Shock 33, 149 (2010).

16. Tattevin, P. et al. Enhanced Indoleamine 2,3-Dioxygenase Activity in Patients with Severe Sepsis and Septic Shock. J. Infect. Dis. 201, 956–966 (2010).

17. Wirthgen, E. & Hoeflich, A. Endotoxin-Induced Tryptophan Degradation along the Kynurenine Pathway: The Role of Indolamine 2,3-Dioxygenase and Aryl Hydrocarbon Receptor-Mediated Immunosuppressive Effects in Endotoxin Tolerance and Cancer and Its Implications for Immunoparalysis. J. Amino Acids 2015, (2015).

18. Litzenburger, U. M. et al. Constitutive IDO expression in human cancer is sustained by an autocrine signaling loop involving IL-6, STAT3 and the AHR. Oncotarget 5, 1038–1051 (2014).

19. Jung, I. D. et al. Blockade of Indoleamine 2,3-Dioxygenase Protects Mice against Lipopolysaccharide-Induced Endotoxin Shock. J. Immunol. 182, 3146–3154 (2009).

20. Mondanelli, G. et al. A Relay Pathway between Arginine and Tryptophan Metabolism Confers Immunosuppressive Properties on Dendritic Cells. Immunity 46, 233–244 (2017).

21. Hayashi, T. et al. 3-Hydroxyanthranilic acid inhibits PDK1 activation and suppresses experimental asthma by inducing T cell apoptosis. Proc. Natl. Acad. Sci. 104, 18619–18624 (2007).

22. Mor, A., Tankiewicz-Kwedlo, A., Krupa, A. & Pawlak, D. Role of Kynurenine Pathway in Oxidative Stress during Neurodegenerative Disorders. Cells 10, 1603 (2021).

23. Taniguchi, T. et al. Indoleamine 2,3-dioxygenase. Kinetic studies on the binding of superoxide anion and molecular oxygen to enzyme. J. Biol. Chem. 254, 3288–3294 (1979).

24. Grant, R. S., Naif, H., Espinosa, M. & Kapoor, V. IDO induction in IFN-gamma activated astroglia: a role in improving cell viability during oxidative stress. Redox Rep. Commun. Free Radic. Res. 5, 101–104 (2000).

25. Goldstein, L. E. et al. 3-Hydroxykynurenine and 3-Hydroxyanthranilic Acid Generate Hydrogen Peroxide and Promote α-Crystallin Cross-Linking by Metal Ion Reduction. Biochemistry 39, 7266–7275 (2000).

26. Pérez-González, A., Alvarez-Idaboy, J. R. & Galano, A. Dual antioxidant/pro-oxidant behavior of the tryptophan metabolite 3-hydroxyanthranilic acid: a theoretical investigation of reaction mechanisms and kinetics. New J. Chem. 41, 3829–3845 (2017).

27. Darlington, L. G. et al. On the Biological Importance of the 3-hydroxyanthranilic Acid: Anthranilic Acid Ratio. Int. J. Tryptophan Res. IJTR 3, 51–59 (2010).

28. Nyhus, K. J., Wilborn, A. T. & Jacobson, E. S. Ferric iron reduction by Cryptococcus neoformans. Infect. Immun. 65, 434–438 (1997).

29. King, N. J. C. & Thomas, S. R. Molecules in focus: Indoleamine 2,3-dioxygenase. Int. J. Biochem. Cell Biol. 39, 2167–2172 (2007).

30. Kim, S. S., Sohn, J. & Lee, S.-J. V. Immunosenescence in Caenorhabditis elegans. Immun. Ageing 19, 56 (2022).

31. Mándi, Y. & Vécsei, L. The kynurenine system and immunoregulation. J. Neural Transm. 119, 197–209 (2012).

32. Cribbs, A. P. & Williams, R. O. Role of the Kynurenine Pathway in Immune-Mediated Inflammation. in Targeting the Broadly Pathogenic Kynurenine Pathway (ed. Mittal, S.) 93–107 (Springer International Publishing, 2015). doi:10.1007/978-3-319-11870-3_7.

33. Stuhr, N. L. & Curran, S. P. Bacterial diets differentially alter lifespan and healthspan trajectories in C. elegans. Commun. Biol. 3, 1–18 (2020).

34. Sutphin, G. L. et al. Caenorhabditis elegans orthologs of human genes differentially expressed with age are enriched for determinants of longevity. Aging Cell 16, 672–682 (2017).

35. Badawy, A. A.-B. & Guillemin, G. The Plasma [Kynurenine]/[Tryptophan] Ratio and Indoleamine 2,3-Dioxygenase: Time for Appraisal. Int. J. Tryptophan Res. IJTR 12, 1178646919868978 (2019).

36. Espejo, L. S. et al. SICKO: Systematic Imaging of Caenorhabditis Killing Organisms. 2023.02.17.529009 Preprint at 10.1101/2023.02.17.529009 (2023).

37. Espejo, L. et al. Long-Term Culture of Individual Caenorhabditis elegans on Solid Media for Longitudinal Fluorescence Monitoring and Aversive Interventions. J. Vis. Exp. JoVE (2022) doi:10.3791/64682.

38. Coburn, C. & Gems, D. The mysterious case of the C. elegans gut granule: death fluorescence, anthranilic acid and the kynurenine pathway. Front. Genet. 4, 151 (2013).

39. Coburn, C. et al. Anthranilate Fluorescence Marks a Calcium-Propagated Necrotic Wave That Promotes Organismal Death in C. elegans. PLOS Biol. 11, e1001613 (2013).

40. Jung, W. H. & Kronstad, J. W. Iron and fungal pathogenesis: a case study with Cryptococcus neoformans. Cell. Microbiol. 10, 277–284 (2008).

41. Zhang, Y., Wang, F. & Zhao, Z. Metabonomics reveals that entomopathogenic nematodes mediate tryptophan metabolites that kill host insects. Front. Microbiol. 13, (2022).

