## Supplemental Figures for "The Emerging Role of 3-Hydroxyanthranilic Acid on *C. elegans* Aging Immune Function"

### Slide 1
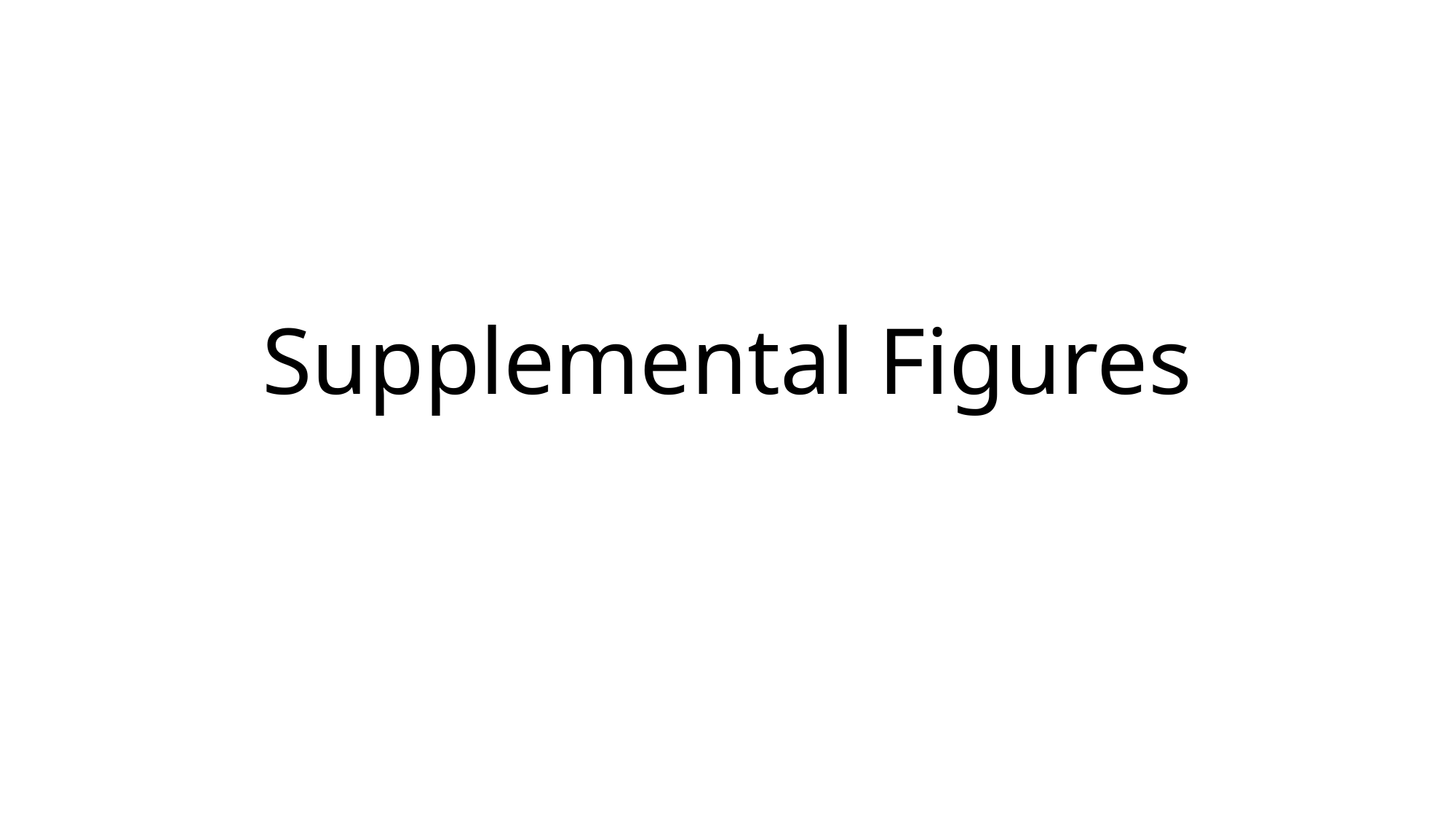

#### Supplemental Figures

### Slide 2
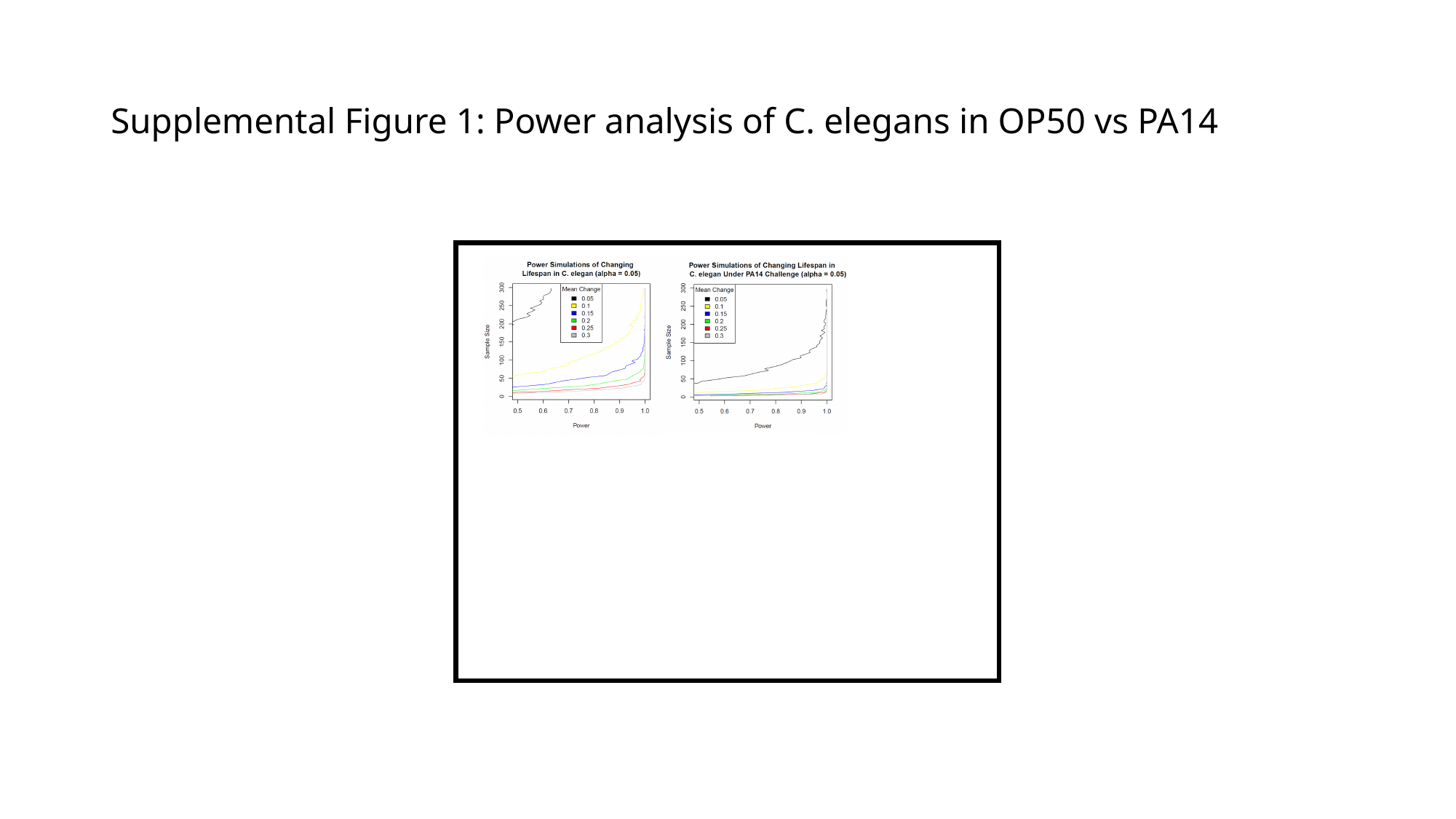

#### Supplemental Figure 1: Power analysis of C. elegans in OP50 vs PA14

### Slide 3
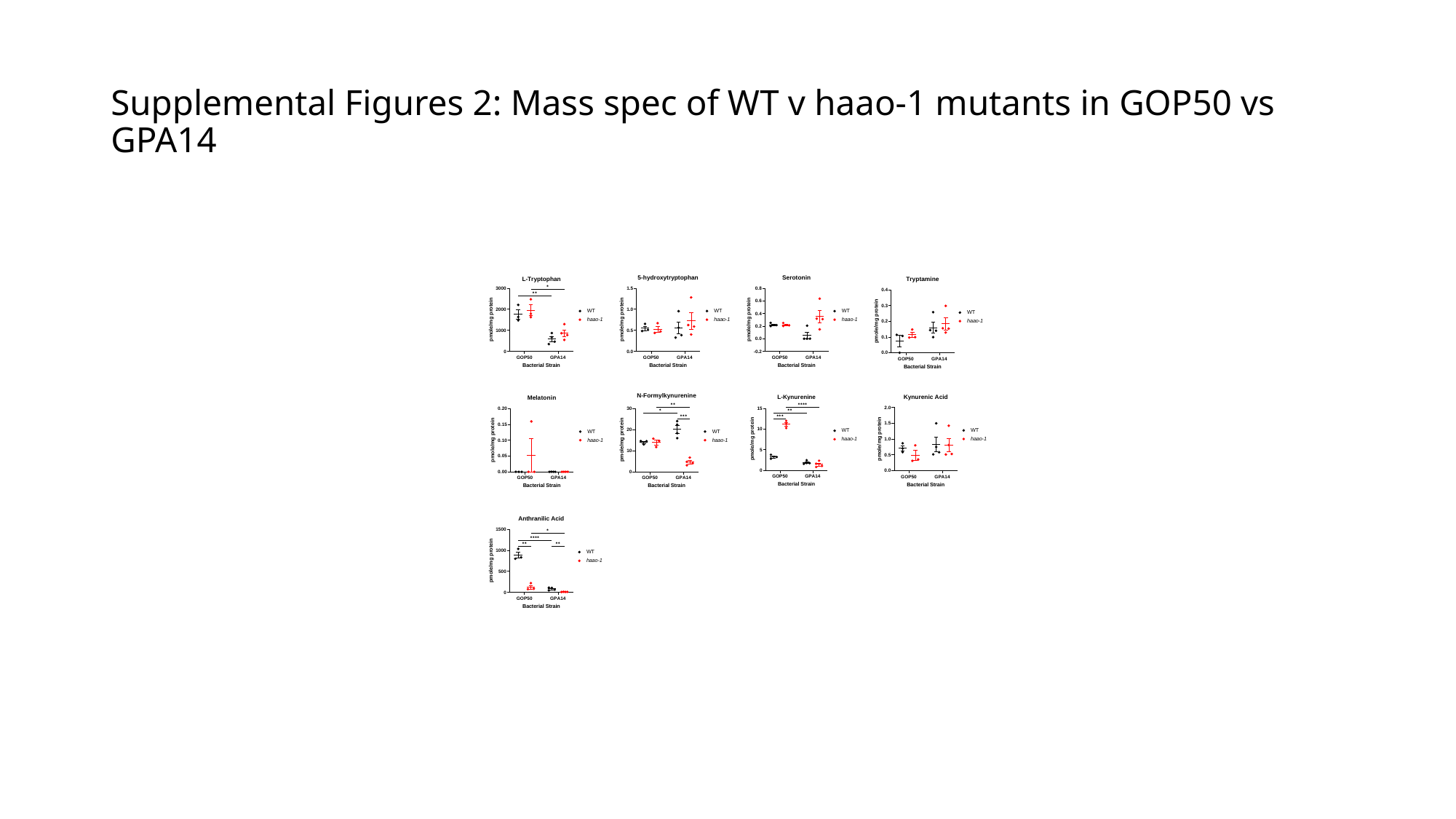

#### Supplemental Figures 2: Mass spec of WT v haao-1 mutants in GOP50 vs GPA14

### Slide 4
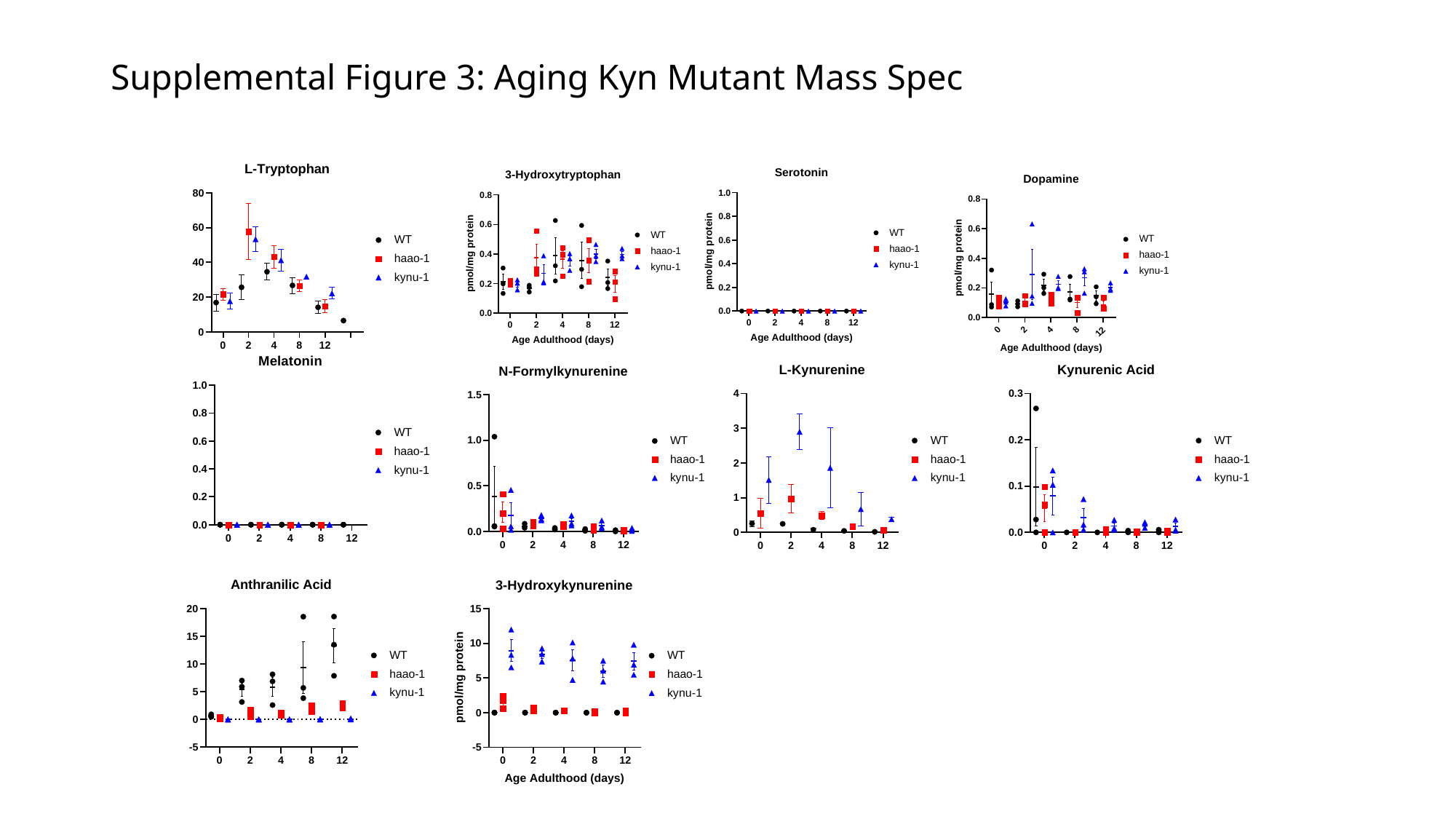

#### Supplemental Figure 3: Aging Kyn Mutant Mass Spec

### Slide 5
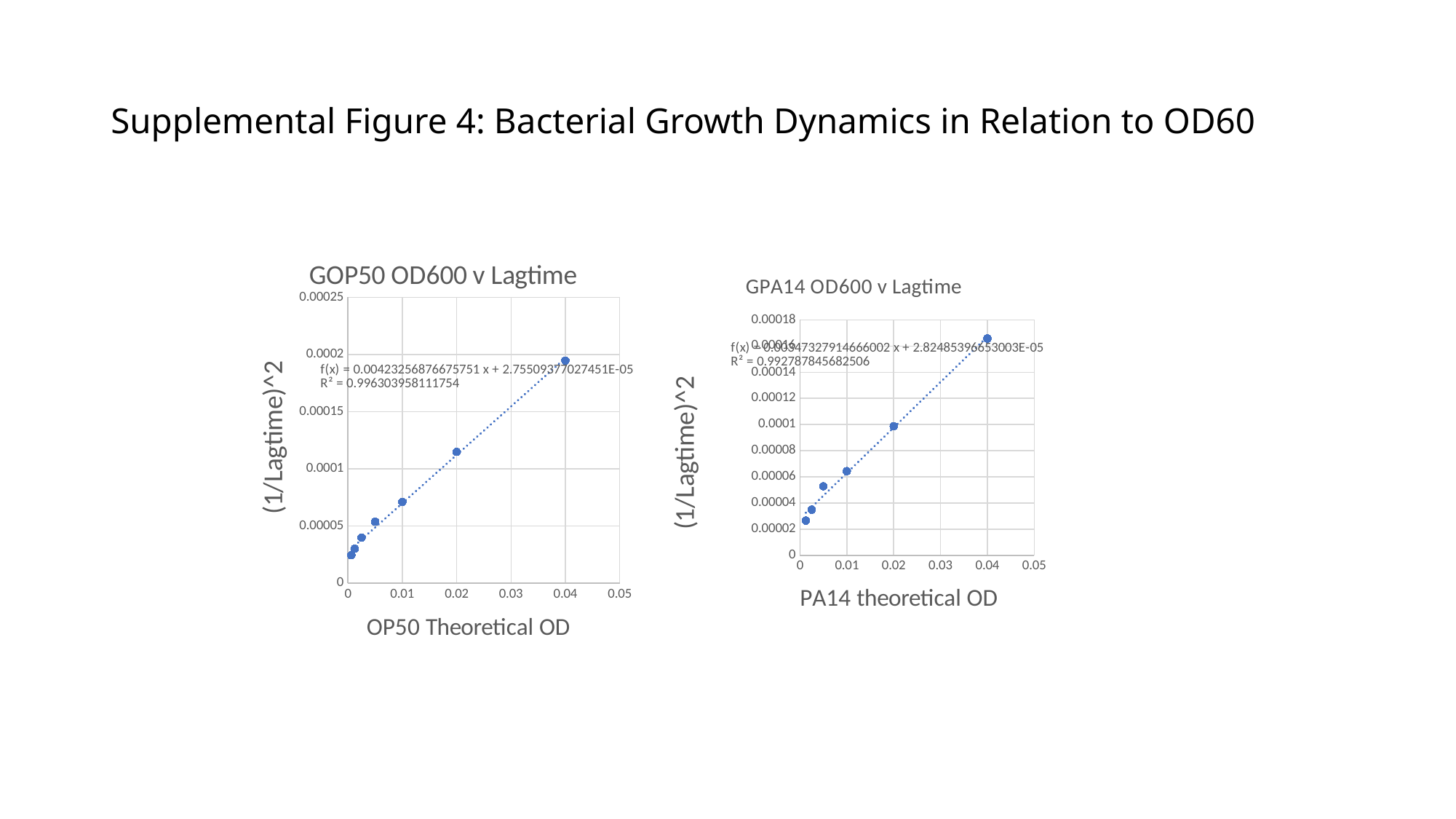

#### Supplemental Figure 4: Bacterial Growth Dynamics in Relation to OD60
###### Chart: GOP50 OD600 v Lagtime
| Category |
|---|
###### Chart: GPA14 OD600 v Lagtime
| Category |
|---|

### Slide 6
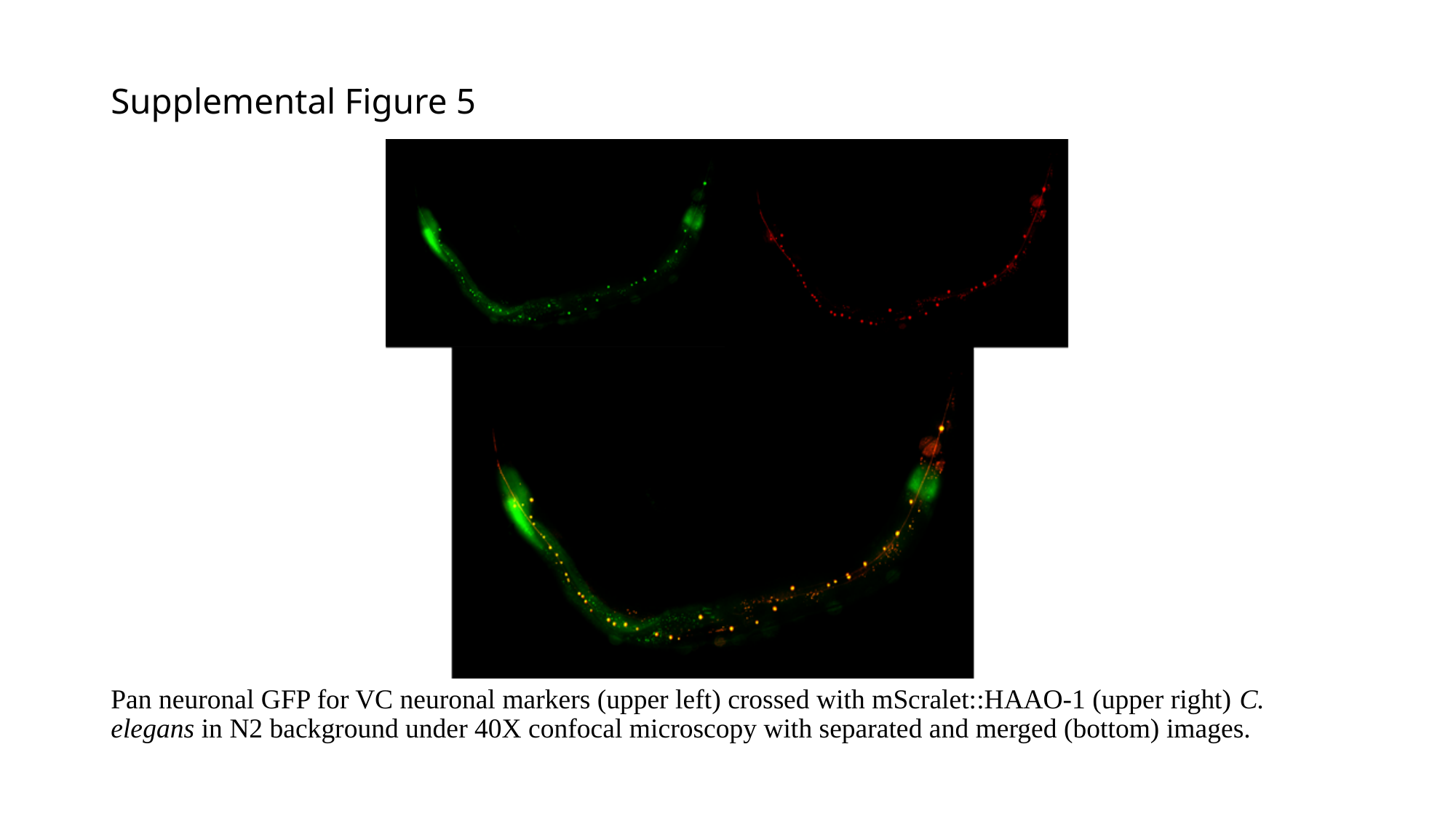

#### Supplemental Figure 5
Pan neuronal GFP for VC neuronal markers (upper left) crossed with mScralet::HAAO-1 (upper right) C. elegans in N2 background under 40X confocal microscopy with separated and merged (bottom) images.

### Slide 7
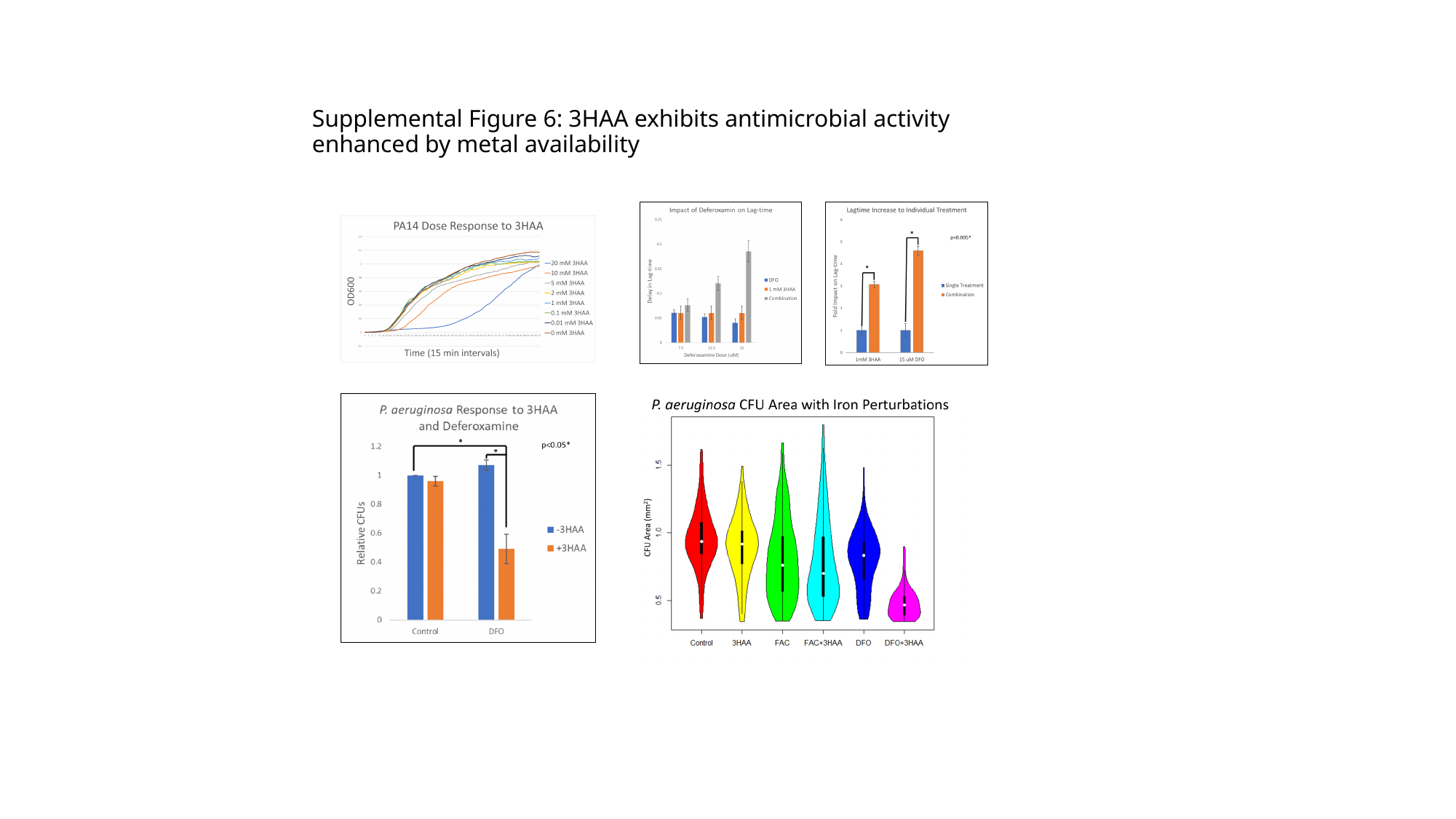

#### Supplemental Figure 6: 3HAA exhibits antimicrobial activity enhanced by metal availability
